# Evaluating Stored Platelet Shape Change Using Imaging Flow Cytometry

**DOI:** 10.1101/2022.05.12.491681

**Authors:** Olga Yakovenko, Tahsin Özpolat, Anastasiia Stratiievska, S. Lawrence Bailey, Jeffrey Miles, Chomkan Usaneerungrueng, Xiaoping Wu, Moritz Stolla

## Abstract

Platelets are routinely stored at room temperature for 5-7 days before transfusion. Stored platelet quality is traditionally assessed by Kunicki’s morphology score. This method requires extensive training, experience, and is highly subjective. Moreover, the number of laboratories familiar with this technique is decreasing. Cold storage of platelets has recently regained interest because of potential advantages such as reduced bacterial growth and preserved function. However, platelets exposed to cold temperatures change uniformly from a discoid to a spherical shape, reducing the morphology score outcomes to a binary “spheres versus discs” during cooling. We developed a simpler, unbiased screening tool to measure temperature-induced platelet shape change using imaging flow cytometry. When reduced to two dimensions, spheres appear circular, while discs are detected on a spectrum from fusiform to circular. We defined circular events as having a transverse axis of > 0.8 of the longitudinal axis and fusiform events ≤ 0.8 of the longitudinal axis. We show that most EGTA-treated platelets and spherical beads are detected in the gate for circular events, while fresh platelets are found mainly in the gate for fusiform events. Using this assay, mouse and human platelets show a temperature and time-dependent, two-dimensional shape change from fusiform to circular, consistent with their three-dimensional change from discs to spheres. The method we describe here is a valuable tool for detecting shape change differences in response to agonists or temperature and will help screen for therapeutic measures to mitigate the cold-induced storage lesion.

**Plain language summary:** *What is the context?:*

- Platelets for transfusion are currently stored for 5-7 days at room temperature, increasing the risk for bacterial growth
- Cold storage reduces the risk for bacterial growth but reduces circulation time
- Stored platelet quality can be assessed by the light microscopy-based Morphology Score, first described in the 1970s
- Downsides of the Morphology Score include subjectivity, extensive training, and reduced availability in platelet laboratories.

*What is new?:*

- In this study, we provide data showing that the Morphology score is reduced to a binary spheres versus discs response in cold-exposed platelets
- We developed an imaging flow cytometry-based approach to quantify platelets’ response to cold based on the two-dimensional projection of the three-dimensional shapes, i.e., fusiform (discoid) versus circular (discoid and spherical)
- We provide validation of this approach in mouse and human platelets
- We provide evidence for increased temperature sensitivity in mouse platelets

*What is the impact?:*

- This study provides an easy and unbiased tool for laboratories working on circumventing the cold-induced storage lesion or documenting spherical shape change in general

## Introduction

Platelets are small discoid blood cells produced by megakaryocytes in the bone marrow at 10^11^/day. Platelets circulate for 7-10 days providing vascular integrity and promoting hemostasis at sites of vascular injury, although a plethora of other functions have been described. Patients with acute hemorrhagia or thrombocytopenia may receive platelet transfusions to prevent and treat bleeding. For this purpose, platelets are currently stored at room temperature for up to 7 days. In the 1960-1970s, platelets were stored at cold temperatures (4 °C), but this approach was abandoned due to decreased *in vivo* circulation time.^1^ Nevertheless, storing platelets at 4 °C has potential advantages, such as the reduced risk of bacterial growth and possibly better function. Overall, these advantages have led to a renewed interest in cold-stored platelets.^2^ Particularly, for patients with acute bleeding during surgery or in a trauma setting, cold-stored platelets could be helpful because long circulation times are not required to promote hemostasis.

Platelet quality during storage is traditionally assessed by a morphology score, described by Kunicki et al. in 1975.^3^ This method requires extensive training and experience. Still, it is one of the few *in vitro* markers that correlates well with *in vivo* recovery and even more so with survival.^3^ In brief, the score is based on four morphologic characteristics, 1) discoid shape, 2) spheroid shape, 3) presence of dendrites (platelets with pseudopodia and dendritic processes), and 4) presence of ballooned platelets (platelets that are unable to maintain the osmotic gradient). One hundred events are counted and multiplied by an arbitrary factor of 4 for discs, 2 for spheres, 1 for dendrites, and 0 for balloons. Hence, a morphology score of 400 indicates platelets with the most desirable morphology (all discs). The current study is aimed at providing an easier, minimalistic and unbiased alternative to the traditional, light microscopy-based morphology score. We report morphology scores from several trials in healthy humans, describe an imaging flow cytometry-based approach, and provide data from validation experiments.

## Materials and Methods

### Reagents

We utilized the following antibodies: CD61-APC, CD42a-FITC (both BD Biosciences, Franklin Lakes, NJ). Low weight heparin enoxaparin (Lovenox, Sanofi-Aventis, Paris, France). Fluoresbrite® YG Microspheres 3.00μm from Polyscience Inc (Warrington, PA, USA).

#### Mouse colony

14-18 weeks old (with body mass about 20-24 g) wild-type C57BL6/J mice from Jackson Laboratory Lab (Bar Harbor, ME) were used in this study. All animal procedures were conducted in accordance with approved institutional IACUC protocols.

### Preparation of platelets

Whole blood was collected by phlebotomy in Na-citrate (3.2%) in plasma from healthy donors. Murine whole blood were collected by retro-orbital bleeding in low molecular weight heparin (Levonox, Sanofi-Aventis, Paris, France). We generated platelet-rich plasma by soft centrifugation (200 g), and platelet-rich plasma was expelled while sparing the buffy coat.

The platelet preparation of the clinical trial morphology score is provided in more detail in the original reports of the studies.^4-6^ Of note, the Morphology Scores have not been previously published.

### Storage Condition

Platelets from mice or human subjects were divided into two equal samples in platelet storage bags. The platelets were either stored at room temperature or at 4 °C. After 1, 24, and 48 hours of storage, the platelets were sampled for the staining for flow cytometry.

### Platelet labeling

We used paraformaldehyde as a fixative known for preserving cell shape. For each time point, 100 mL of 3% paraformaldehyde (PFA) solution (1% final) was added directly to the cell medium to avoid any perturbation in cell shape. After 10 minutes of incubation with PFA, the cells were washed twice with PBS and labeled with CD61 – APC and CD42a – FITC for human platelets. After incubation at room temperature in the dark for 10 minutes, the samples were resuspended in 400 mL of PBS.

### Imaging Flow Cytometry

Samples were analyzed by using Amnis Imagestream Mk II (Luminextcorp, Austin, TX). Images of cells were then acquired with the Amnis Image Stream with the 60X magnitude objective using the bright field channel, the SSC channel, FITC (channel 2) and APC (Channel 11) by the INSPIRE software for acquisition. IDEAS 6.2 software (Luminexcorp, Austin, TX) were used for analysis. First, we identified focused platelets based on bright-field images by the levels higher than 60 on gradient root mean (GRM) square channel 1 (M01) feature to analyze platelet morphology. We isolated platelets based on CD61 positivity (we gated for CD61 positive cells based on 1e^3^-2e^5^ MFI). In the third step, we defined single platelets by using the Area and the Aspect ratio features. Lastly, we used aspect ratio (circular events with a transverse axis of > 0.5 of the longitudinal axis, and fusiform events with ≤ 0.5 of the longitudinal axis) to analyze different morphologies of stored platelets (Figure 2 A). The boundaries of the gates were defined based on the shape of circular versus fusiform dot plots. This was further refined and confirmed with image controls.

### Healthy human subjects research

The Western Institutional Review Board approved the research (WIRB), and all human participants gave written informed consent. The study was conducted following the Declaration of Helsinki.

### Statistical analysis

We reported the results as mean ± standard error of the mean and assessed for statistical significance by paired Student t-test with correction for multiple comparisons. A P value of ≤ 0.05 was considered significant.

## Results

One hallmark of platelets exposed to cold temperature is the change from discoid shape to spherical shape, based on tubulin disintegration and actin barbed end-capping.^7^ Indeed, we have included the Kunicki morphology score in previous cold storage and room temperature storage studies and found it to be uniformly 200 (all spheres), at least for the first 15 days of storage (Figure 1). In contrast, storing platelets at 22 °C increases the score number and variability (Figure 1). To avoid using this complex and subjective method for evaluating morphologic change in cold-stored platelets, we utilized imaging flow cytometry. We based our gating strategy on the premise that spheres can only be detected as circles when reduced to two dimensions, but disks can be seen as fusiform (i.e. spindle-shaped) events or circles and on a spectrum between both shapes. Therefore, we first aimed at selecting a gating strategy that allowed us to differentiate between fusiform and circular events. We used the ratio of the longitudinal to the transverse axis. Circular events were defined as having a transverse axis of > 0.5, while fusiform (spindle-shaped) events were defined as having a transverse axis of ≤ 0.5 of the longitudinal axis. The final gates were based on aspect ratio and intensity threshold (Figure 2A-D). In addition to the bright-field analysis, we added fluorochrome-labeled antibodies to platelet surface markers to exclude the attachment of occasional leukocytes or RBCs to platelets (Figure 2B, Figure 3, A, B). After 48 hours of room temperature storage, we saw a mixed distribution between circular and fusiform events, but 48 hours of cold storage led to mostly circular events (Figure 2E, F).

**Figure 1:**
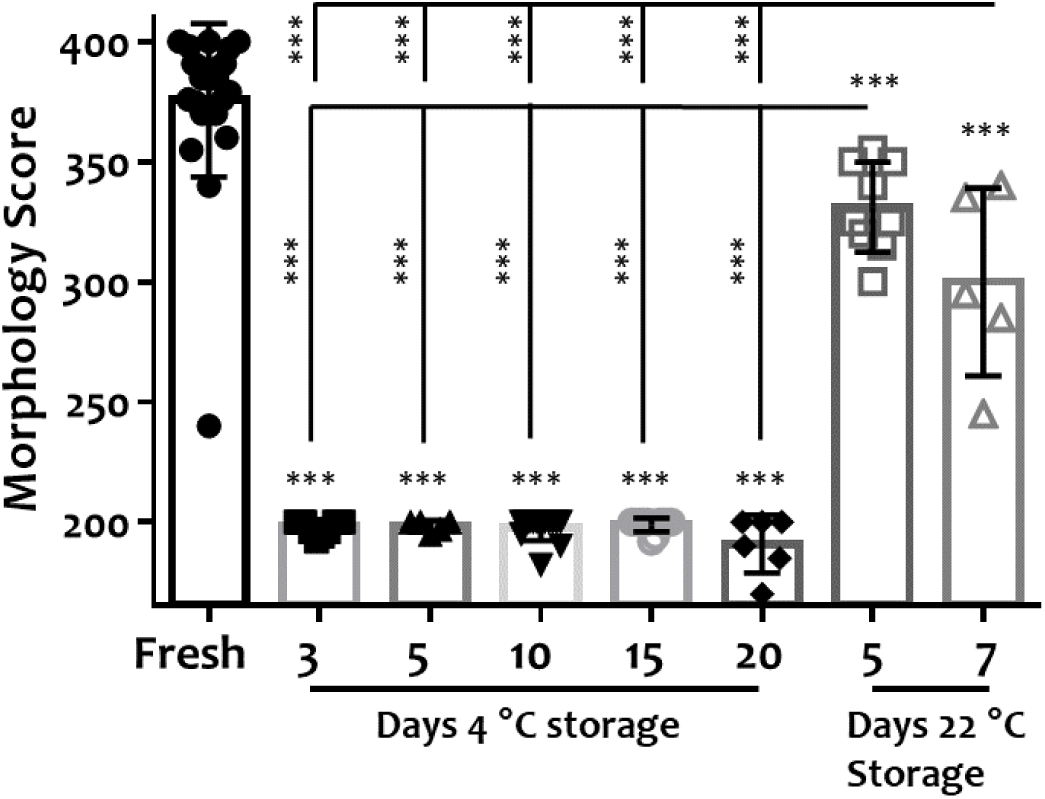
Morphology scores for fresh platelets, 3, 5, 10, 15, 20 day cold-stored, and 5, 7 room temperature-stored platelets. (Fresh (black circles) n=25, Cold-stored platelets, duration: 3 days (black squares, n=12), 5 days (black upward triangles, n=5), 10 days (black downward triangle, n=14), 15 days (grey circles, n=11), 20 days (black diamonds, n=6), RT-stored platelets, duration 5 days (grey squares, n=9), 7 days (grey squares, n=5). Data show as individual data points, mean, and ± SD. ***p ≤ 0.001.

**Figure 2:**
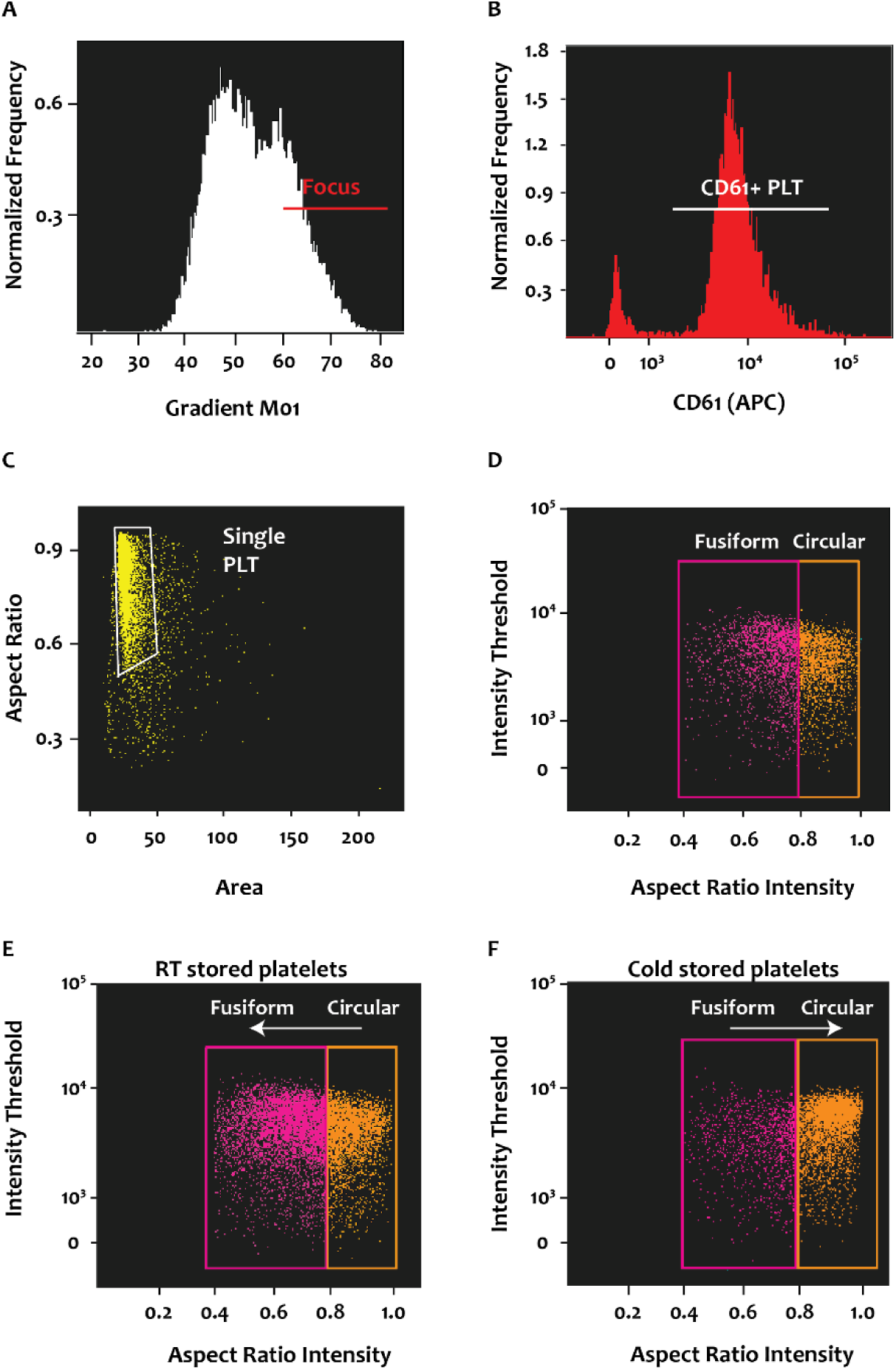
Quantification of cold-induced platelet shape change by imaging flow cytometry. **(A-D)** Gating Strategy to differentiate between fusiform and circular platelets. Circular events were defined as having a transverse axis of >0.5 of the longitudinal axis and the percentage of circular (orange gate) and fusiform platelets (pink gate) was determined **(D)**. After 48h of RT storage, 31% were in the gate for fusiform platelets, and 64% were in the gate for circular platelets **(E)**. After 48h of cold storage, 96% of platelets were in the gate for circular platelets and only 6 % were in the gate for fusiform platelets **(F)**.

**Figure 3:**
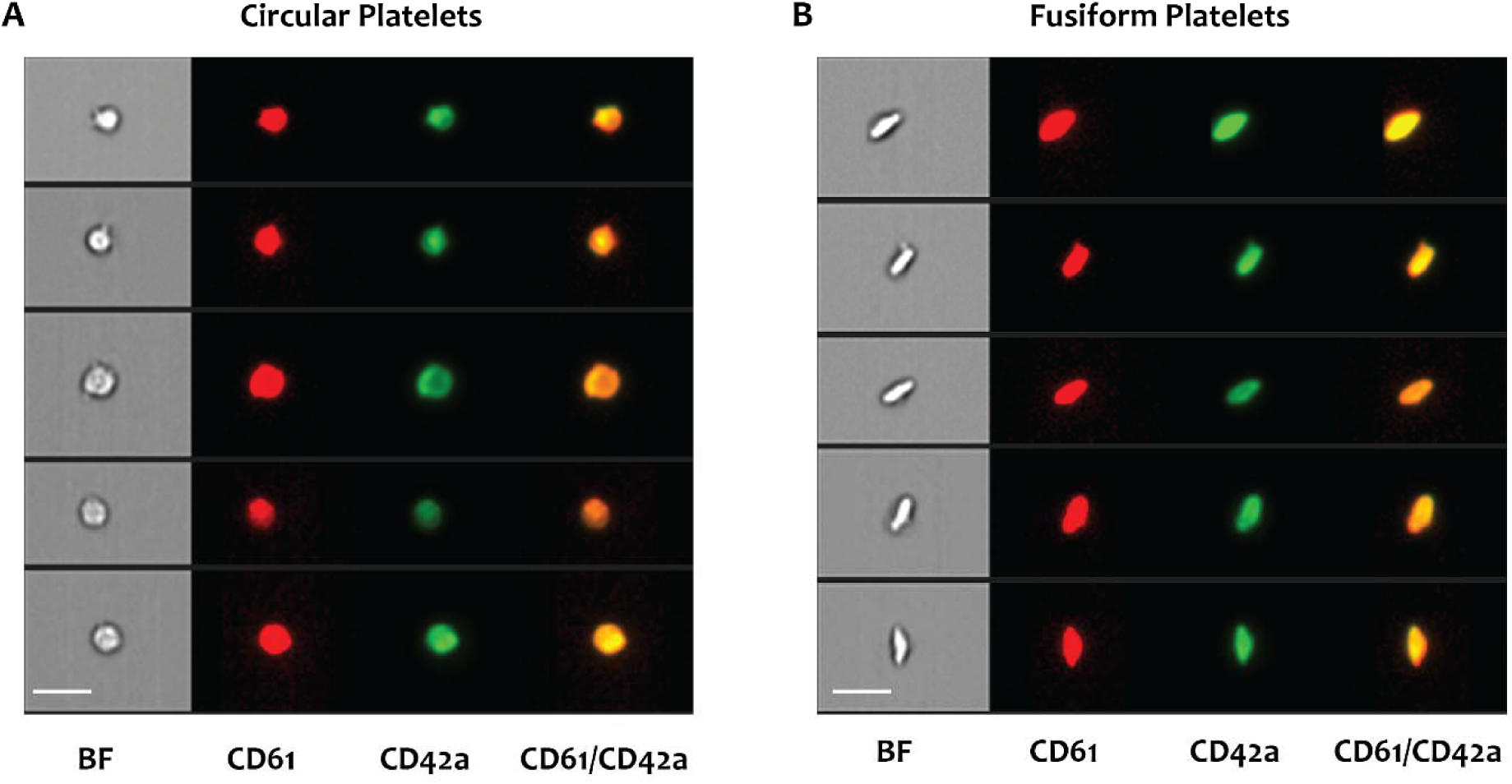
Representative samples for fusiform and circular events. Five random events are shown for circular **(A)** and fusiform **(B)** platelets.

The calcium chelating reagents EDTA and EGTA induce a spherical shape change in platelets.^8,9^ To validate the methodology and test whether we can differentiate an entirely spherical platelet population from a mostly discoid population, we treated fresh platelets with EGTA and compared their appearance to untreated, fresh platelets. Indeed, we found EGTA-treated platelets predominantly in the circular gate (Figure 3A), while fresh control platelets were mostly gated as fusiform events (Figure 4A). Similarly, spherical beads were almost exclusively identified in the circular gate (Figure 4A). We exposed mouse and human platelets to temperatures between 37 °C and 4 °C for 60 minutes and subsequently examined them in our assay. In both species, lowering the temperature led to a decline in fusiform events and increased circular events, consistent with a transition from fusiform to spherical shape (Figure 4B). The lower the temperature, the more likely platelets were to be detected in the gate for circular events. Of note, mouse platelets show a slightly higher crossover point with more circular than fusiform events than human platelets (Figure 3B, lower panel), indicating increased temperature sensitivity. Even when exposed to 4 °C, one hour was insufficient to result in a significantly higher circular population in human platelets, while mouse platelets showed significantly more circular events at 10 °C and below. To further study the time-dependency of cold-induced shape change in human platelets, we exposed human platelets to one, 24, and 48 hours to room temperature or 4 °C in commercially available storage bags. Even after prolonged storage, room temperature led to an even distribution between circular and fusiform events (Figure 4C, upper panel). In contrast, cold exposure led to a trend for more circular events after 24 hours of cold storage (Figure 4C, lower panel), which further increased after 48 hours.

**Figure 4:**
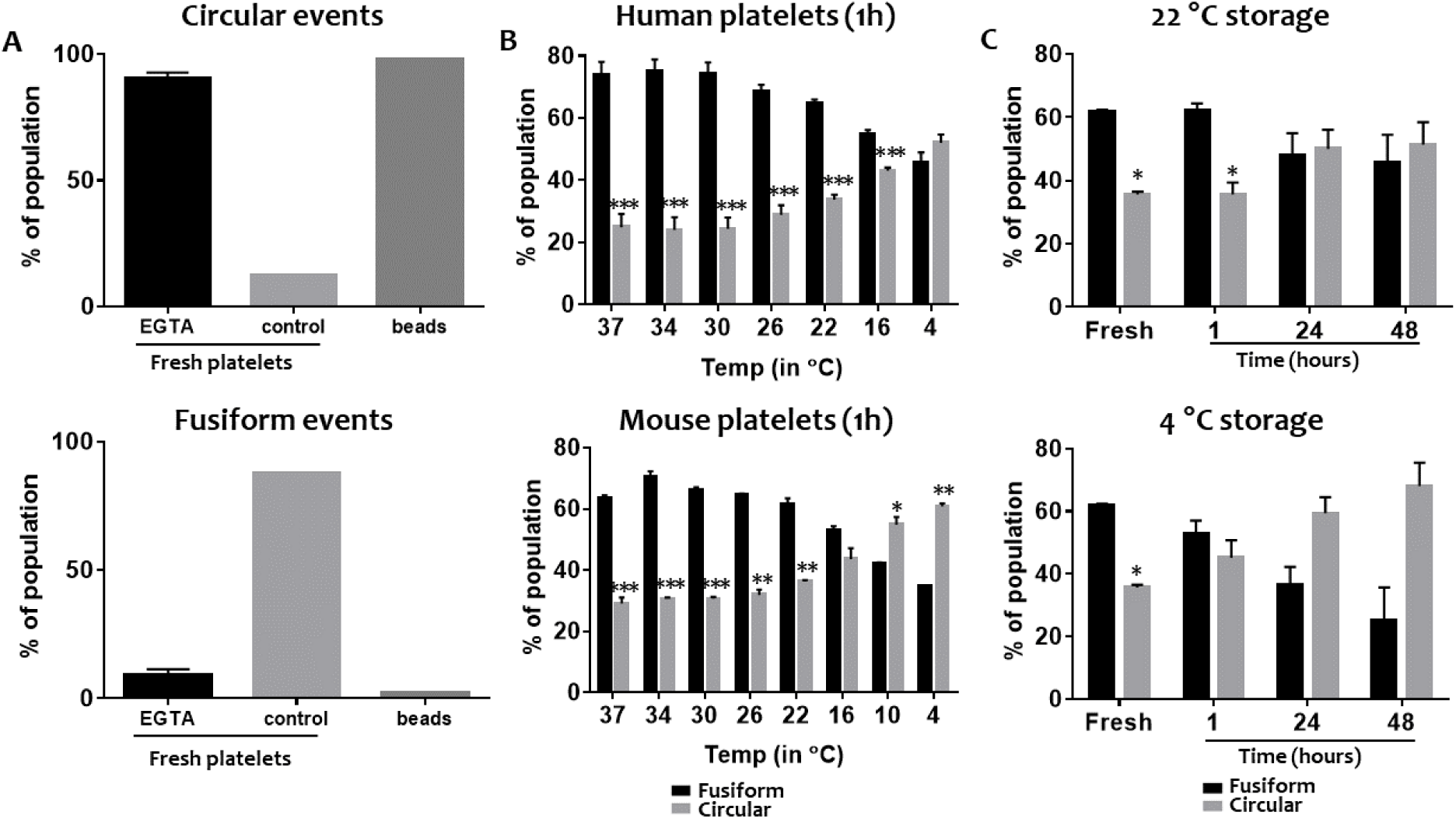
**(A)** To validate the methodology, platelets were left untreated (discoid control) or treated with EGTA to induce spherical shape change. Spherical beads were included as a non-platelet spherical control. The percentage of circular events is shown on the upper panel and fusiform events on the lower panel. **(B)** Human platelets (upper panel) and mouse platelets (lower panel) were exposed to temperatures from 37 °C to 4 °C for one hour, fixed, and tested by imaging flow cytometry. N=4, ***p ≤ 0.001, **p ≤ 0.01, *p ≤ 0.05 **(C)** Human platelets were stored at room temperature (22 °C, upper panel), or 4 °C (lower panel) for up to 48 hours and tested as baseline (fresh), and after one, 24, and 48 hours, N=2, *p ≤ 0.05.

## Discussion

Platelets are routinely stored for transfusion to bleeding patients or patients at risk for bleeding. The transfusion community recently re-discovered an interest in cold storage of platelets, intending to transfuse safer and potentially more efficacious platelets. Current research projects aim at preventing shape change and, ultimately, longer circulation times. Although initial studies had suggested that shape change itself is not the primary reason for clearance^10^, shape change still represents the hallmark of the cold-stored platelet storage lesion and the prevention thereof, a primary research goal of the transfusion community.^11^ The platelet quality of room temperature-stored platelets is historically assessed by the Kunicki morphology score. Our data show that after extended periods of cold storage up to 15 days, the score is uniformly resulted as 200 (all spheres). Hence, we did not note any correlation with *in vivo* recovery and survival (data not shown). Other tools based on the discoid shape of “healthy” platelets have similar problems, e.g., the “swirling phenomenon” based on the movement and orientation of discoid platelets in plasma is uniformly absent in cold-stored platelets (data not shown).

Moreover, obtaining reliable results with the morphology score requires extensive training, and even after completion of training, the high level of subjectivity inevitably leads to inter-operator and intra-operator variability. Because light microscopy used for the morphology score is not designed to assess three-dimensional objects either, obtaining the morphology score routinely involves physical manipulation of the cell suspension (i.e. slight movement of the coverslip) to characterize the three-dimensional shape of platelets. We developed an imaging flow cytometry-based tool to quantify the degree of cold-induced shape change based on the two-dimensional projection of three-dimensional objects. Our approach omits subjectivity and represents an unbiased tool to quantify the degree of shape change for future research and clinical studies. This initial validation study shows how uniformly spherical suspension and discoid cells appear in the gates designed to detect circular and fusiform events. Staining with platelet markers further decreased the chance of including false positive (non-platelet) events. We found mouse platelets to be more temperature-sensitive than human platelets. To our knowledge, this has not been described in the literature before. However, mouse platelets are known to be hyperreactive compared to human platelets, and the increased sensitivity to cold temperature may be a reflection of that, as many of the consequences of cold exposure resemble agonist stimulation.^11,12^ In summary, we developed a standardized screening tool to assess and quantify storage condition-induced platelet shape change. Our assay will help screening for novel drugs and approaches to circumvent the unwanted side effects of cold storage, including shape change.

## Author contribution

O.Y. performed experiments, analyzed data, and co-wrote a first darft of the manuscript, T.O. performed experiments, analzyed data, providede critical technical help, and helped writing the manuscript, A.S. analyzed data and helped writing the mansucript, S.L.B, J.M., and C.U. all performed experiments and analyzed data, X.W. provided critical technical help and helped writing the manuscript, M.S. designed the study, analyzed data, and co-wrote a first draft of the manuscript

## Acknowledgments

The authors would like to thank Renetta Stevens and Tena Petersen for their administrative support. Funding sources: NIH (1R01HL153072-01), DoD (W81XWH-12-1-0441, EDMS 5570), American Society of Hematology Scholar Award.

